# Rehumanizing the homeless: Altered BOLD response following contact with an extreme outgroup

**DOI:** 10.1101/462671

**Authors:** P. A. Kirk, A. O. Cohen, W. Sinnott-Armstrong, L.T. Harris

**Affiliations:** Institute of Cognitive Neuroscience, University College London, UK; NYU Department of Psychology, New York University, USA; Kenan Institute of Ethics, Duke University, USA; Department of Experimental Psychology, University College London, UK

**Keywords:** Dehumanization, Contact, Social Cognition, fMRI

## Abstract

Dehumanized perception strips people of their humanity by ignoring their mental states. Evidence has accumulated to suggest many individuals do not spontaneously attribute mental states towards certain social groups, such as the homeless, and drug-addicted (Fiske et al., 2002, Harris & Fiske, 2006; 2007; 2011). These groups tend to elicit differential BOLD signal within areas associated with social cognition. To investigate the versatility of this response, two experiments were conducted: a mixed design study (20 participants); and a repeated-measures design (11 participants). These investigated the malleability of social cognition following a contact intervention with the homeless. Both experiments had participants make emotional judgements toward eight different social groups whilst under fMRI. The two studies found mixed evidence, demonstrating altered response to homeless people in regions such as the mPFC, Insula, IPL, and MTG following social contact. This lends some support to the use of contact as an effective intervention.

---

People spontaneously attribute mental states to other people (Young & Saxe, 2009). Making inferences about others’ minds is at the very foundation of social cognition, informing moral judgements (Young, Cushman, Hauser, & Saxe, 2007), and is a prerequisite for empathy (Shamay-Tsoory, 2011). Yet, when viewing certain populations, people do not extend this fundamental component of social communication, failing to infer their mental states (Harris & Fiske, 2006; 2007; 2011). Although these dehumanized perceptions are often discussed in relation to violence and war (Grossman, 1996, p. 188), it does not only occur in such extreme contexts; ordinary people’s disengagement of social cognition towards the homeless is frequent (see Harris & Fiske, 2009 for a review), and such everyday dehumanization can result in apathy towards providing aid (Cuddy, Rock, & Norton, 2007). On a socio-political scale, some theorize dehumanized perception encompasses cyclical, self-fulfilling consequences, and results in support for aggressive policies towards said social groups (Kteily, & Bruneau, 2017). Destructive and forceful legal action taken towards the homeless (Pasha-Robinson, 2018) exemplifies this. Finding a way to re-humanize these social groups, particularly the homeless, thus remains a priority.

## A Framework of Dehumanization

The continuum model of impression formation (Fiske, & Neuberg, 1990; Fiske, Lin, & Neuberg, 1999) remains a useful framework from which to view social cognition. In addition to the idea that person perception occurs along a continuum, a pertinent concept of the model is that humans make rapid assessments or categorizations based on easily accessible features, such as race, gender, and age. Only after this, if one has sufficient information and are so motivated, do individual characteristics influence person perception. The model suggests schema-triggered affect in response to the social group moderates this process. Detailing this affective influence, the Stereotype Content Model (SCM; Fiske, Cuddy, Glick, & Xu, 2002) outlines how stereotyped warmth (positive/negative intent), and competence (high/low capability) can predict perceivers’ emotional response. These models laid a foundation for the dehumanization literature, demonstrating that those in the low warmth, low competence group (low-low, extreme outgroup) fail to elicit spontaneous social cognition (in United States (US) samples).

## Studies of Dehumanization

An early fMRI study (Harris & Fiske, 2006) investigated behavioral and brain responses to the homeless, and drug-addicted, both traditionally dehumanized social groups in the low-low outgroup. The results indicate pictures of these groups as evoking feelings of disgust, a non-social emotion, as opposed to other groups that evoked the social emotions of pride, pity, and envy. The authors suggested these feelings of disgust to the low-low groups explained the increased activity in the amygdala and anterior insula. Moreover, while targets outside the low-low groups evoked a significant increase in the social cognition brain network, especially in the medial prefrontal cortex (mPFC), the homeless, and drug-addicted targets did not, indicating dehumanized perception.

Subsequent research (Harris & Fiske, 2007) used the same paradigm, but asked participants to determine a targets’ age or vegetable preference, rather than report evoked emotions. This study replicated the reduced engagement of the social cognition brain network to the low-low groups. However, when asked to infer vegetable preference (compared to the age of the targets), participants increased activity in the social cognition brain network, including regions of the mPFC, inferior parietal lobe, middle temporal gyrus, and insula. This suggested that moving beyond stereotypes and inferring mental content information re-engages social cognition. Evidence outside the dehumanization literature supports this – promoting broad, abstract mindsets increases judgements solely on stereotype information (McCrea, Wider, & Myers, 2012).

A two-part behavioral study (Cameron, Harris, & Payne, 2016) further sheds light on these results. Participants were expecting an emotionally distressing video of a homeless person. Those who reported greater feelings of anticipated exhaustion typically attributed less mental states to the homeless individual. The second experiment manipulated how mentally exhausting the participants were told the video would be. Their results revealed stigma – exhaustion – mental state inference interaction: mental state inferences decreased when there was greater anticipated exhaustion, though this was primarily for stigmatized targets (drug users). Both experiments lend support to the notion that dehumanized perception can arise as a proactive emotion regulation strategy to protect oneself from imagined negative mental experiences. This claim is consistent with data showing that participants who scored high in empathy dehumanize more, and that nurses who humanize their patients are more likely to hold symptoms of burnout (Vaes, and Muratore, 2012). However, the latter may not always be the case, especially in people who work with the homeless (see Ferris et al., 2016).

## Intergroup Contact

While the previous literature has laid the foundation for the neural correlates of dehumanized perception, no fMRI study to date has investigated experimentally manipulating this response as a product of an established, real-world intervention. Contact provides us with a promising method for investigating this. Since intergroup contact’s original conception (Allport, 1954), numerous studies have supported its use for reducing intergroup bias. Meta-analyses of the research reveals contact correlates with a reduction in prejudice (Pettigrew & Tropp, 2006), and is mediated by anxiety, perspective taking/empathy, and outgroup information (Pettrigrew & Tropp, 2008).

To date, there have been a handful of neuroimaging studies including prior contact as a correlational variable. The first of its kind, one study (Cloutier, Li, & Correll, 2013) had participants making perceptual judgements of other-race faces whilst undergoing fMRI. The researchers showed a tendency for lower elicitation of amygdala activity as a product of childhood intergroup contact. Brain imaging research in the same year (Telzer, Humphrey, Shapiro, & Tottenham, 2013) also looked at white participants viewing black faces. Results showed attenuation of the amygdala relates to intergroup contact during adolescent development. However, the authors were not concerned with extreme outgroups, and the former study’s task dissuaded participants from attributing mental states, thus it is unsurprising altered BOLD response was not seen in areas recruited for social cognition. It is also the case that dehumanized perception is not a prejudice response centered on the amygdala. Therefore, a motivating question for this current study is whether effects of contact generalize beyond prejudice (a primarily affective response in contrast to social cognition, which is cognitive). Furthermore, though the researchers established a link between social contact and brain response, they did not manipulate intergroup contact. For the current study the critical question is then – can contact as an intervention extend to extreme outgroups, and is this reflected by a dehumanized/humanized brain response?

The aim of the present research is to investigate the extent to which real-world contact may alter brain responses to an extreme outgroup. We hypothesized that dehumanized perception would be observed in participants when viewing images of homeless people, as indexed by the social cognition network. Following an intervention of contact with this group however, said activity would increase for the homeless, but not another extreme outgroup. Furthermore, it was hypothesized that images of extreme outgroups would evoke feelings of disgust, as recorded behaviorally, and indexed by the amygdala and insula. We predicated a reduction in these affective reactions following the contact intervention.

### Experiment 1: Mixed design

The goal of the experiment 1 was to provide initial evidence that the social cognition brain network response can be modified through social interaction, and to identify brain regions beyond those previously noted, that may be regulating observed patterns of activation.

## Methods

### Participants

We recruited 20 (5 male, 15 female) participants with a mean age of 27.8 years from the Duke-UNC Brain Imaging Analysis Center and Duke Center for Cognitive Neuroscience subject pool and compensated with payment. All participants spoke English as a first language; were right-handed; reported no abnormal neurological conditions, head trauma, or brain lesions; had normal or corrected-to-normal vision; and provided informed consent. 9 participants identified as ‘White/Caucasian’, 6 as ‘Asian’, 2 as ‘African American’, 2 as ‘Hispanic’, and 1 as ‘Indian’. In regards to socioeconomic status, 11 participants identified as ‘low-middle’, 3 as ‘high-middle,’ and 6 as ‘high’.

### Materials

Images of individuals from 8 different social groups appeared in the experiment, totaling 240 different photographs (120 for each scanning session). These were either humanized targets (American, Business, Disabled, Old, Rich, and Student) or dehumanized targets (Drug-addicted, and Homeless). All stimuli had been pre-tested showing a humanized/dehumanized response and above chance for being rated as inducing feelings of pride, envy, pity, or disgust (Harris & Fiske, 2011). We controlled for low-level visual features such as eye gaze. A picture of a landscape marked the start of every run.

### Procedure

We scanned participants on two separate occasions, 5-7 days apart. The overall procedure consisted of the first scanning session (pre-intervention), intervention, and the second scanning session (post-intervention). In each fMRI session we had participants viewing pictures of the various social group members and giving emotional judgement ratings (described below). Before each scanning session, participants gained familiarity with the paradigm outside of the scanner. In between the two scanning sessions, we took participants to the Urban Ministries of Durham soup kitchen. During the subsequent social interactions with homeless people, participants had a list of eight questions regarding the homeless person’s preferences. We instructed participants to have a conversation with the homeless people with the goal of answering the eight questions. We outlined that the interaction could last up to 30 minutes, though how long they stayed was of their choosing. Participants were timed and asked questions regarding the quality of the interaction; specifically to rate their enjoyment of the interaction and familiarity with the homeless person.

Both scanning sessions employed the paradigm used in a previous experiment (Harris & Fiske, 2006), bar the durations noted. Participants viewed a picture of a target; a fixation-cross appeared (jittered 2, 4, 6, or 8 seconds); participants judged what emotion they associated with the picture (pride, envy, pity, or disgust; 2 seconds); and a final crosshair (jittered 2, 4, 6, or 8 seconds) appeared. Participants had 120 trials in each scanning session, spread across six runs, with a randomized presentation order.

### fMRI Acquisition Parameters

We conducted fMRI scanning at the Duke-UNC Brain Imaging Analysis Center using a 3 Tesla GE Signa Excite. A computer projecting to a screen mounted at the rear of the scanner bore presented stimuli, which the participants view while supine through a series of mirrors. A Resonance Technology fiber optic response box (four buttons) recorded the participants’ ratings. Following a brief localizer scan, a T1-wieghted anatomical scan (256 x 256 matrix, 176 1-mm sagittal slices) ran at the beginning of each scanning session, lasting ~12 minutes. T2*-weighted scans were collected using a single-shot gradient echo EPI sequence of the whole brain (39 slices, 3×3×3 mm voxels, TR = 2000ms, TE = 25ms, FOV = 192cm, Flip Angle = 75°, bandwidth = 4340 Hz/px; Echo spacing = .29ms).

### fMRI preprocessing

We performed preprocessing and analysis using the BrainVoyager QX software package. Preprocessing consisted of correction of slice acquisition order; 3D rigid-body motion correction; voxel-wise linear detrending across time; and temporal band-pass filtering to remove low-and high-frequency (scanner- and physiology-related) noise. Correcting for distortions in the EPI image consisted of a simple affine transformation. We coregisted functional images to the structural images, then to standard space (Talairach, & Tournoux, 1988), and applied an 8-mm full-width/half-maximum filter.

### fMRI Analysis

We conducted data analysis using the general linear model (GLM) available in the BrainVoyager QX software package. We first computed a GLM for the pre-interventions scans, collapsed across all subjects, focusing on the 2s periods during which images of humanized vs dehumanized targets were presented. Two separate GLMs were computed for post-intervention scans — one for the video condition, and one of the interaction condition.

Whole-brain analysis was conducted to isolate regions of interest (ROIs) based on the relevant social cognition areas isolated from the contrast of humanized > dehumanized targets in the baseline scan. Given the number and types of stimuli presented (2 dehumanized categories, 6 humanized catergories), a 3:1 deviant cell contrast best examined the data. Additionally, we had a priori anatomically defined ROIs. We compared the beta values from the pre-intervention scans to the post-intervention scans to obtain mean percent signal change values. We then performed Random effects analyses on all imaging data. Finally, repeated measures analyses of the variance (ANOVAs) and correlations with scores from behavioral measures (time, enjoyment rating, familiarity rating) were performed on the beta weights extracted from all ROIs.

## Results

### Behavioral Data

Participants chose to spend significantly more time in the interaction condition than the video condition, which was found using an independent-samples t-test to compare the two conditions, *t* (18) = 4.39, *p* < .00035 (Figure 1). Moreover, time was found to positively correlate with participant’s enjoyment ratings (*r* = 0.656, *p* = 0.002, *n* = 20; Figure 2).

**Figure 1.**
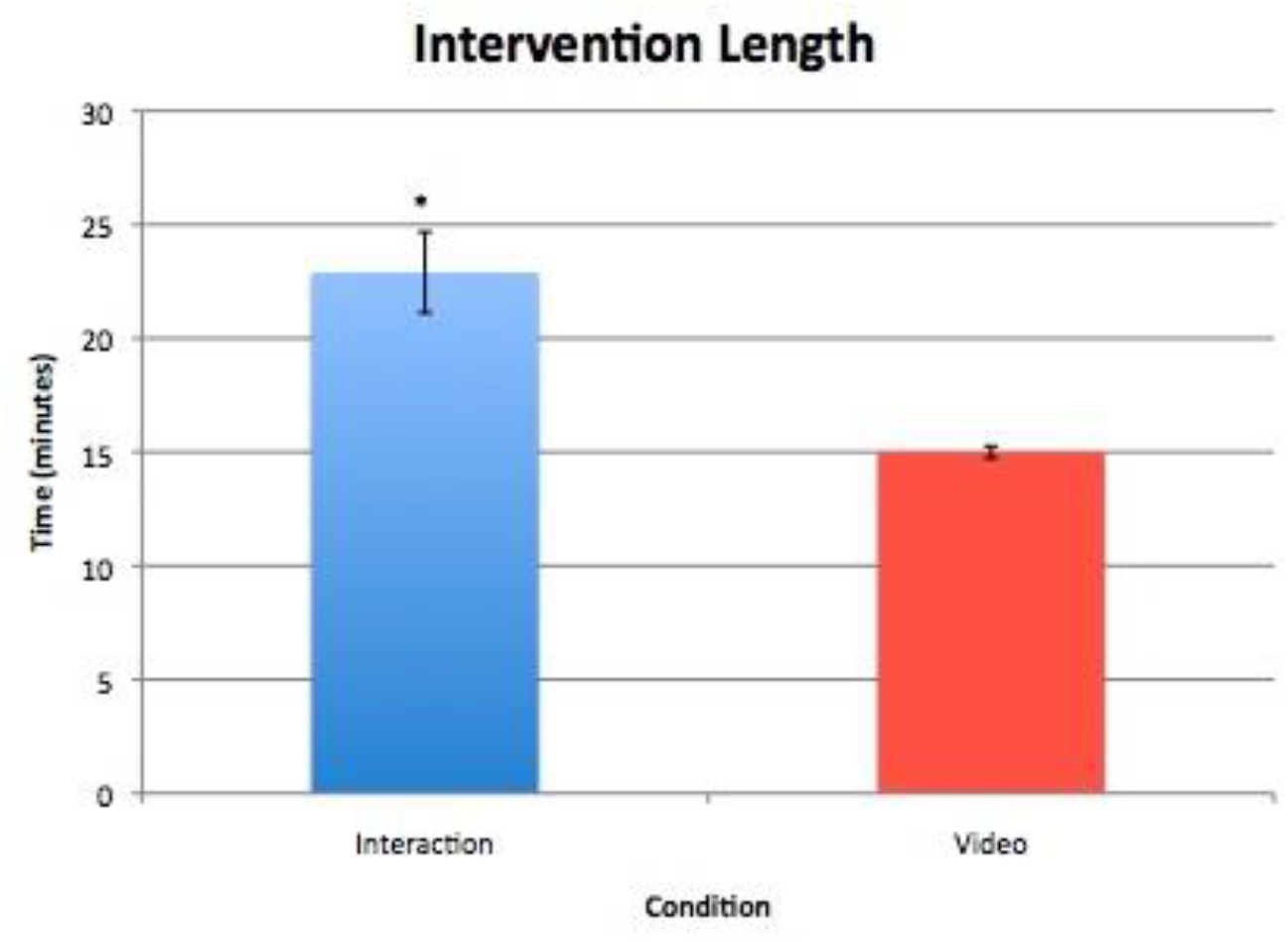
Time spent in intervention. * = significant at p < .00035.

**Figure 2.**
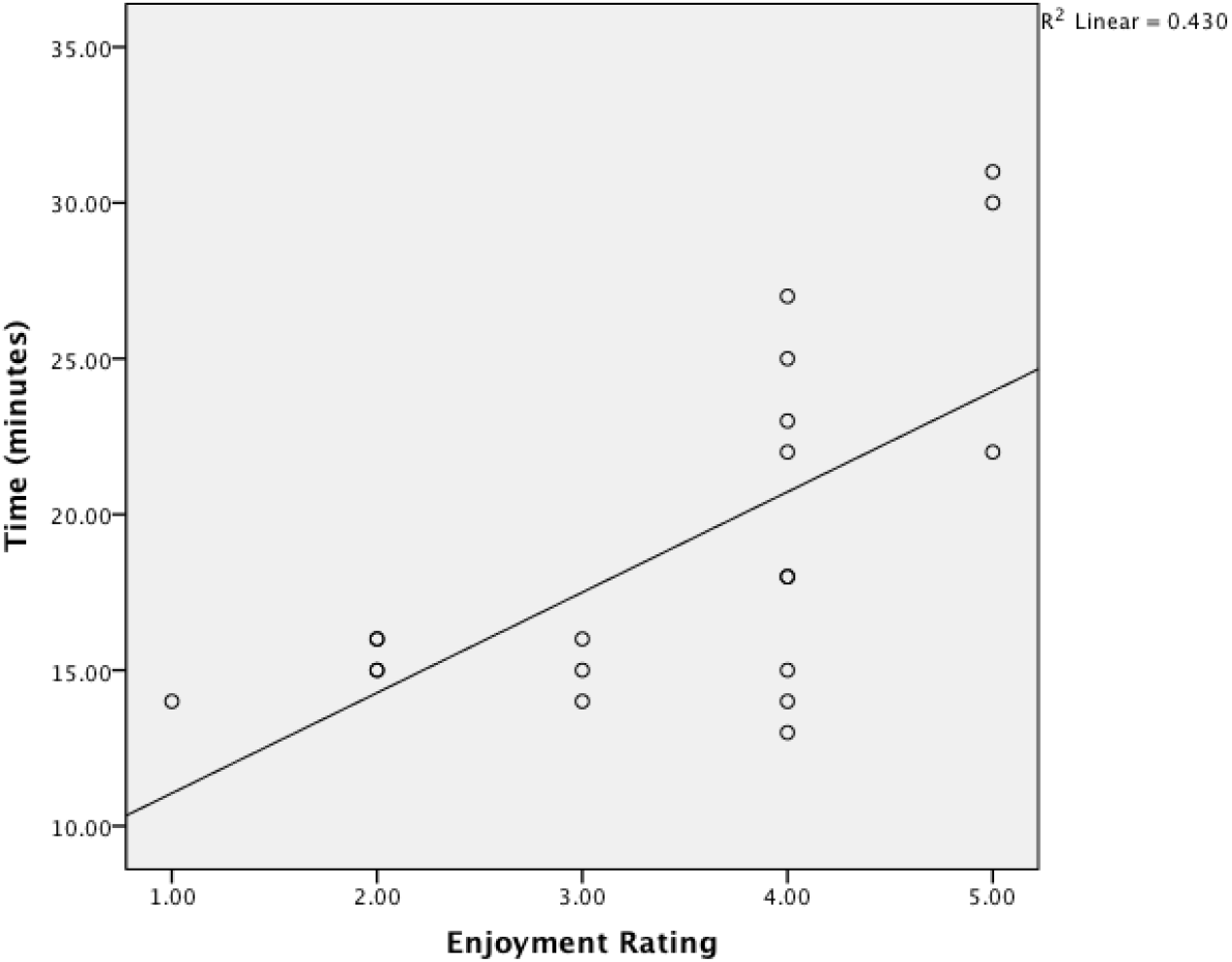
A scatterplot of time spent in interaction v. subjective enjoyment rating with regression line fitted.

We next examined the post-intervention ratings between subject conditions (see Figure 3). A repeated measures ANOVA compared enjoyment and familiarity ratings within and between subjects in each condition. We found a significant main effect for rating scale type (enjoyment or familiarity), *F* (1, 18) = 4.95, *p* < .039, *partial η^2^* = 0.216, *Ω* = 0.557, qualified by a significant interaction effect between scale type and condition, *F* (1, 18) = 12.66, *p* < .002, partial *η^2^* = 0.412, *Ω* = 0.920. Two independent samples t-tests made post-hoc comparisons between conditions. The first t-test indicated no significant difference in familiarity ratings between conditions. The second t-test indicated a significant difference in the enjoyment ratings between conditions *t* (18) = 4.98, *p* < 0.0001.

**Figure 3.**
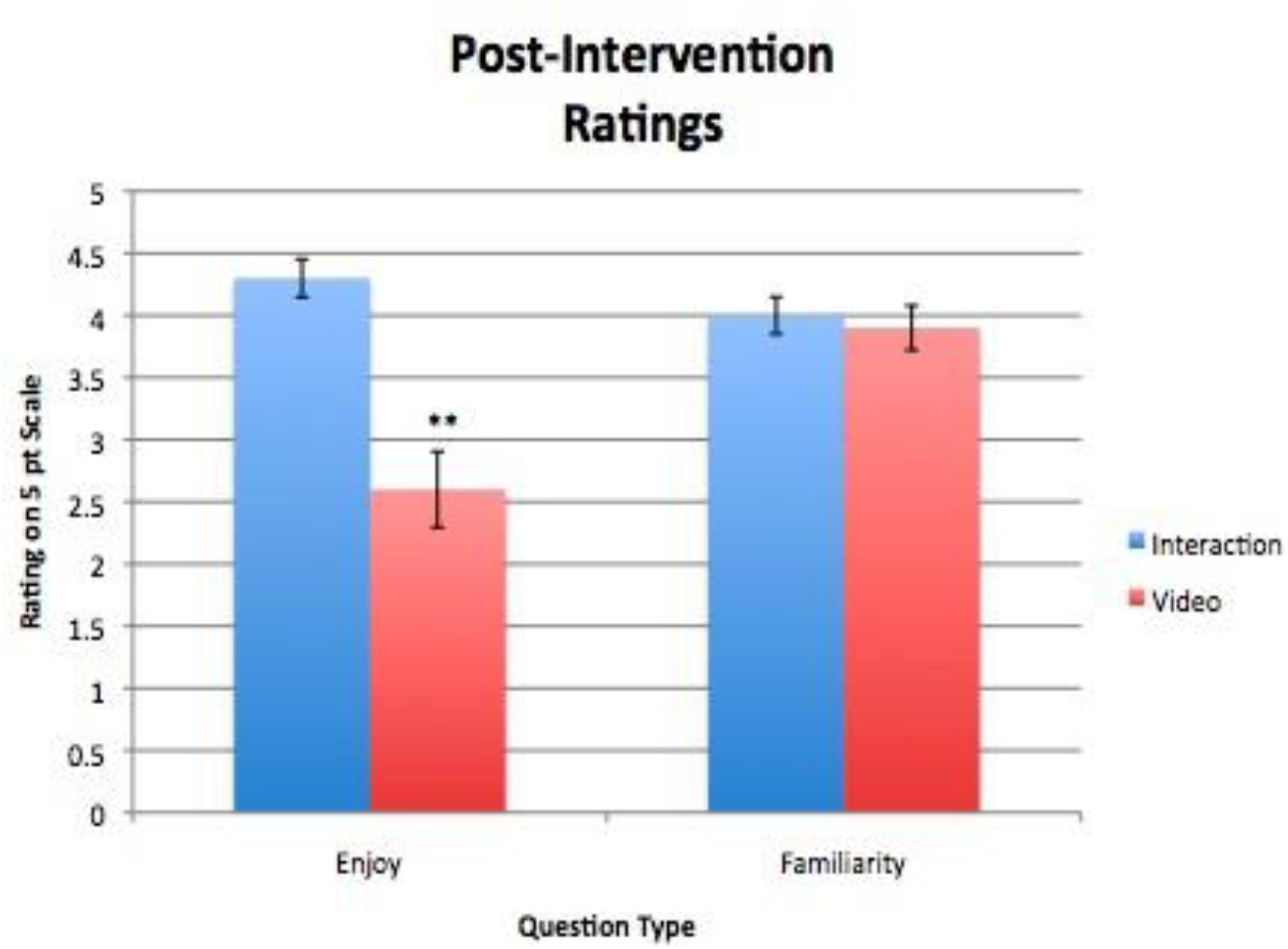
Post-intervention ratings of enjoyment/familiarity. ** = significant for *p*< 9.76E^-5^.

### fMRI Data

#### Whole-Brain Contrast

We conducted a whole-brain analysis in the pre-intervention scan (collapsed across all participants) using a 3:1 contrast with humanized targets > dehumanized targets to isolate any discrepancies in social cognition processes. Consistent with previous findings and the hypothesis, we observed greater activity in the ventral mPFC, *t* (19) = 3.94, *p* < .001, 35 voxels (Figure 4) for humanized over dehumanized targets.

**Figure 4.**
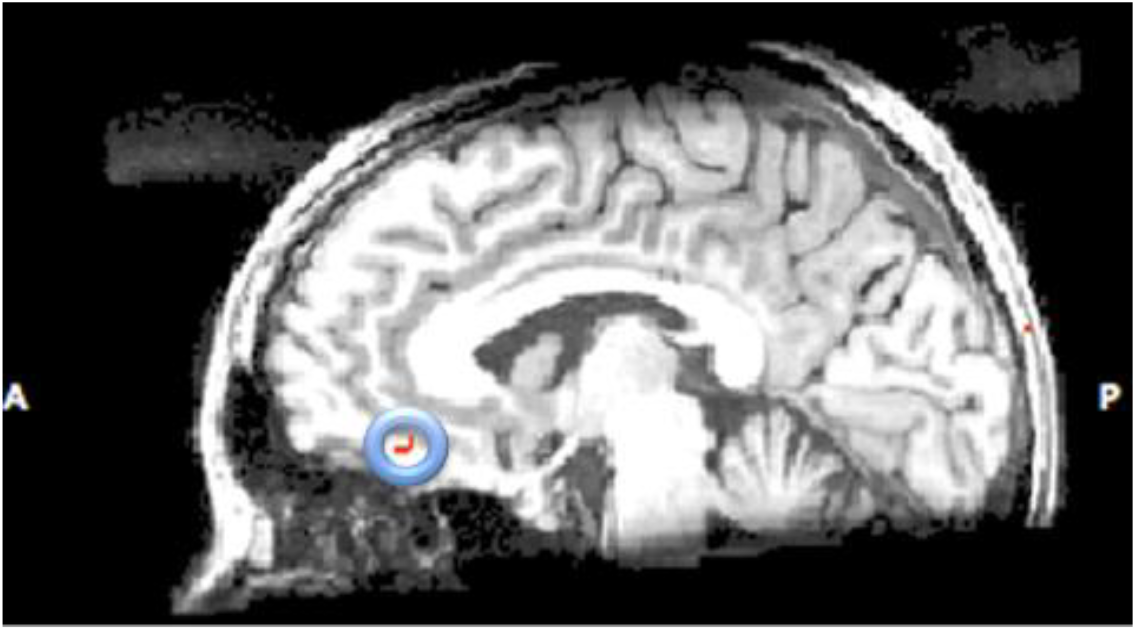
Saggital view of mPFC activation in humanized > dehumanized contrast.

#### Regions of Interest (ROI) Analysis

Our follow-up ROI analysis looked at the mean percent signal change in the functionally defined mPFC region post-intervention, for both conditions, in response to images of homeless individuals. As a point of relevant comparison, we included changes in response to drug-addicted targets. Our one-way repeated measures ANOVA compared mean percent signal change in the interaction and video conditions. Results were trending in the right direction, with increased mean percent signal change in the experimental condition and a decrease in the control. There was no difference between conditions in the mean percent signal change in response to drug-addicted targets, suggesting that the experimental manipulation may or may not have resulted in some form of neural change specifically in response to the homeless (Figure 5). Additionally, signal change in the mPFC for the experimental group (viewing the homeless) positively correlates with time spent in interaction (r = 0.667, p = 0.035, n = 10; Figure 6) and familiarity ratings (r = 0.656, p = 0.039, n = 10; Figure 7), with no such effects found in the control group.

**Figure 5.**
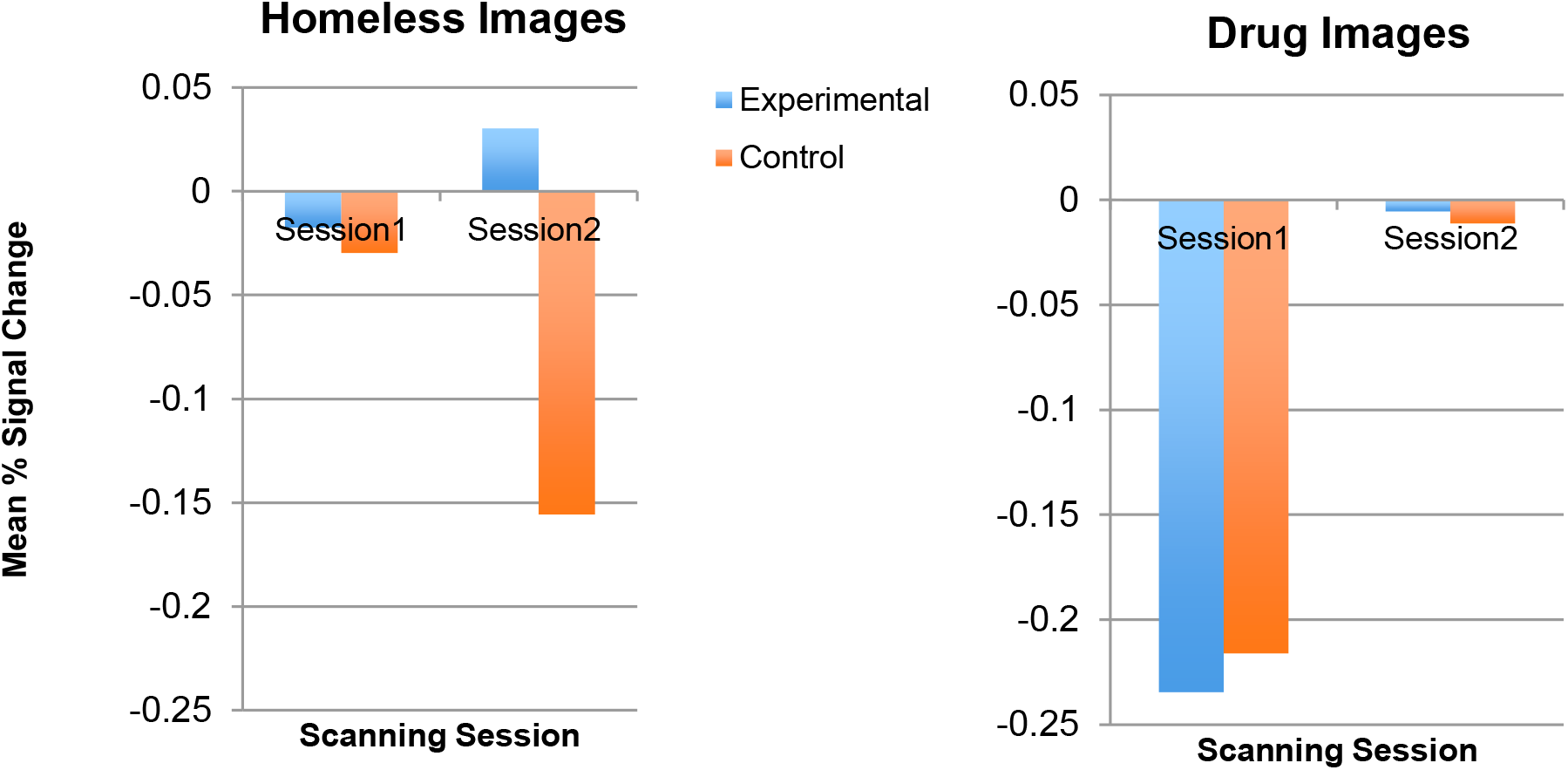
Changes in BOLD signal activation calculated from mPFC region betas

**Figure 6.**
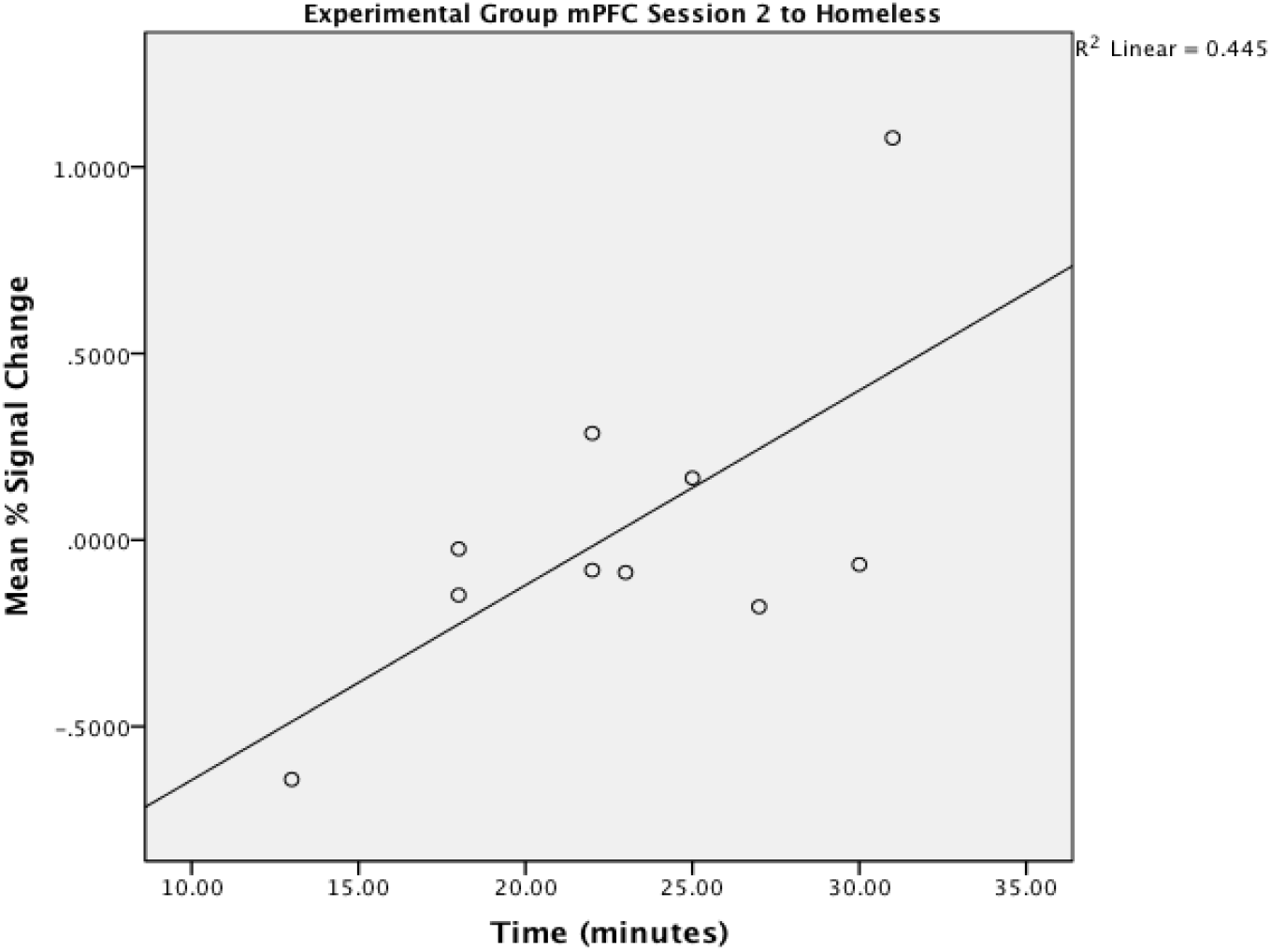
Scatterplot of change in mPFC beta weights vs. time spent in interaction with regression line fitted.

**Figure 7.**
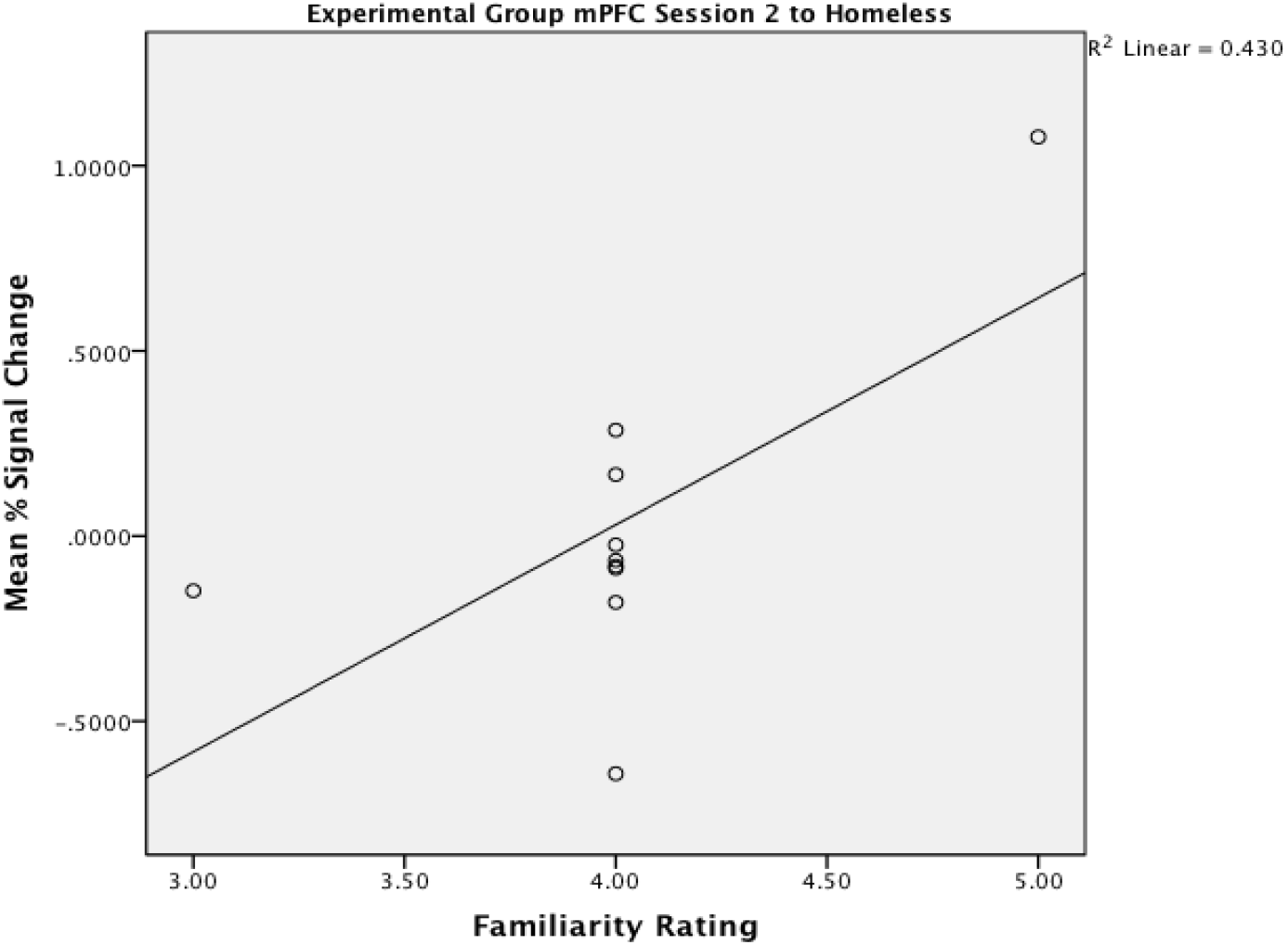
Scatterplot of change in mPFC beta weights vs. familiarity ratings with regression line fitted.

Our subsequent repeated measures ANOVAs compared the mean percent signal change in the interaction and video conditions in anatomically defined bilateral amygdala and bilateral insula, yielding no significant results between conditions. However, we saw several correlations between BOLD signal and behavioral measures. Left amygdala beta weights in response to drug-addicted targets was found to negatively correlate with time spent in the intervention (*r* = −0.469, *p* = 0.037; Figure 8). Additionally, we found marginal (*p* < .10) negative correlations between bilateral insula to drug-addicted targets at session one and participant’s enjoyment ratings of the intervention.

**Figure 8.**
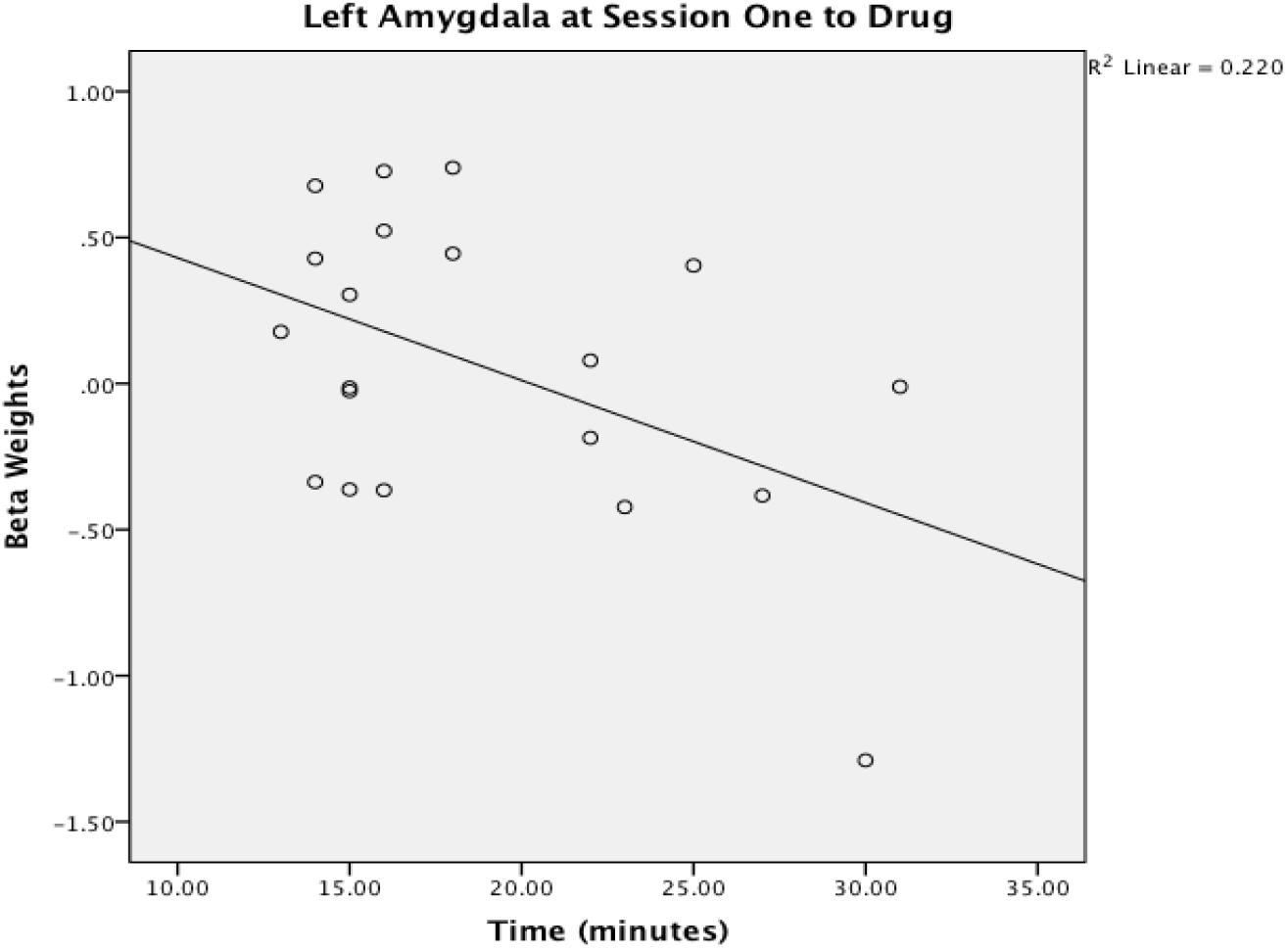
Scatterplot of change in beta weights in the left amygdala vs. time spent in intervention with regression line fitted.

Session two beta values averaged within each condition also revealed correlations. We found left amygdala signal change of the control group in response to business people positively correlated with participant’s enjoyment ratings of the video (*r* = 0.696, *p* = 0.025; Figure 9). Right insula activity in the experimental group in response to Americans showed a marginal positive correlation with enjoyment rating of the interaction. In the control group, left insula activity in response to homeless people showed a marginal negative correlation with time spent watching the video.

**Figure 9.**
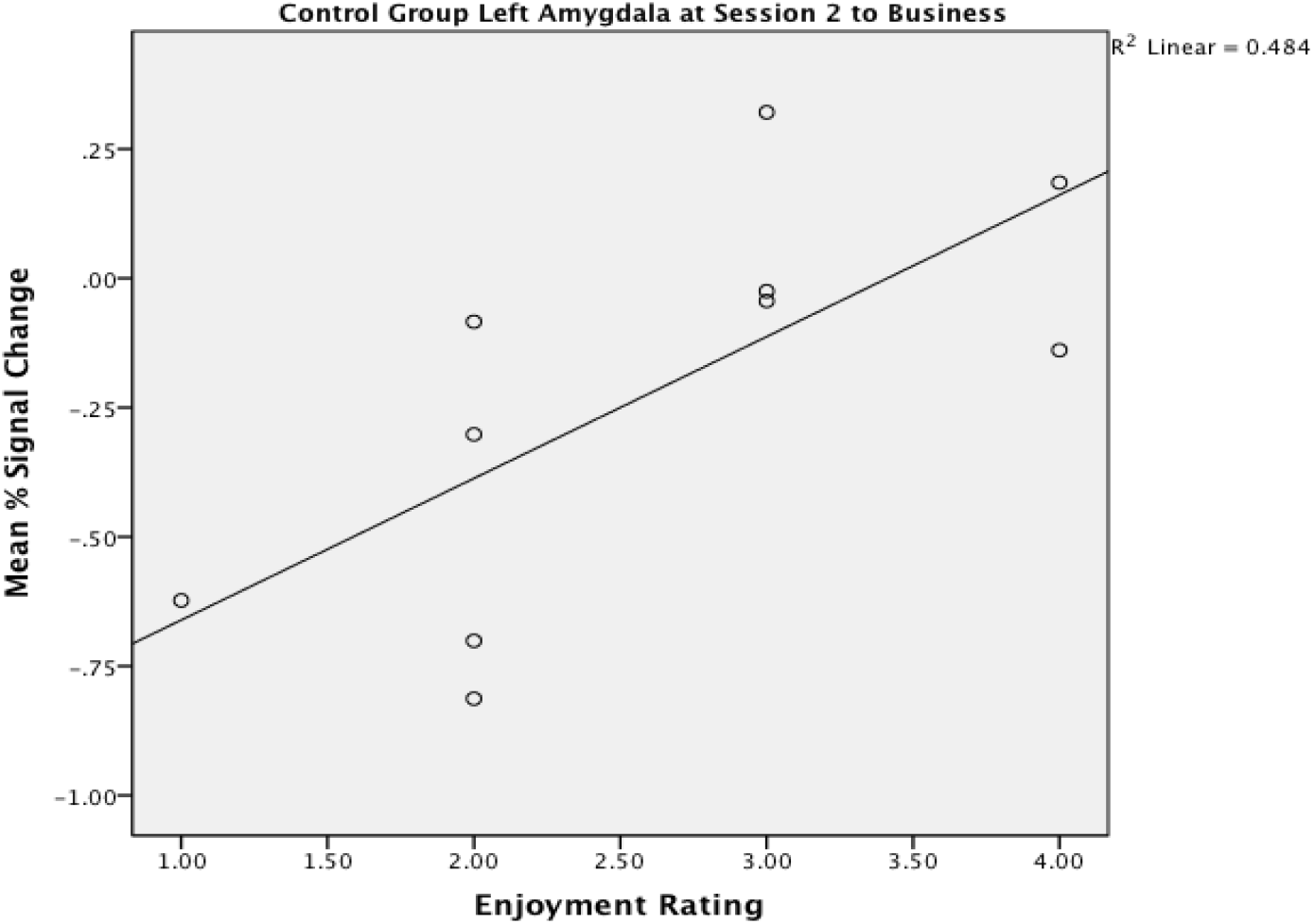
A scatterplot of change in beta weights in the left amygdala vs. enjoyment rating with regression line fitted

### Summary of results

Our results showed decreased mPFC activity in response to dehumanized targets at session, replicating previous findings. The interaction-intervention condition was trending towards engaging the same localized region of mPFC in response to the homeless—but not drug addict targets— more than the video condition at session two. How the amygdala and insula modulate this response remains unclear, and explain these signals via feelings of fear, or disgust. mPFC activity post-intervention significantly correlated with how long participants engaged in the interaction, as well as familiarity ratings of the homeless. Furthermore, left amygdala activity pre-intervention predicted time spent in the intervention.

#### Experiment 2: Replicating the rehumanization effect

With the first experiment leaving multiple questions regarding the mechanisms underlying contact, and the relevant affective responses, the second experiment aimed to shed light on these results with a repeated-measures design using a further 11 participants.

## Methods

### Participants

We recruited 11 participants (3 male, 6 females, 1 unknown) with a mean age of 27.60 years (*SD* = 15.21 years) for the study using the same criteria outlined in experiment 1. 7 participants identified as ‘White/Caucasian’, 2 as ‘Asian’, and 1 as ‘African American’ (demographic information was missing for one participant). In regards to socioeconomic status, 5 participants identified as ‘middle’, 4 as ‘low-middle’, and 1 as ‘high’. 6 participants identified as ‘liberal’, 3 as ‘moderate’, and 1 as ‘conservative’.

### Stimuli, Task, and Procedure

We replicated the stimuli, procedure, paradigm, and fMRI acquisition parameters from the first experiment in a repeated-measures format (no control condition).

### fMRI Preprocessing

We conducted preprocessing and analysis in MATLAB (The Mathworks, Natick, MA) using Statistical Parametric Mapping 12 (Wellcome Trust Centre for Neuroimaging, London, UK). For DICOM-NIFTI conversion, we made use of MRIcron’s DCM2NII converter (Rordern, Karnath, & Bonilha, 2007). Preprocessing took place in the following order: realignment and unwarping; coregistration; segmentation; normalization using segmentation deformation field; and smoothing of the functional images with a full-width at half maximum 8-mm Gaussian kernel.

### fMRI Analysis

We conducted first level analysis at the within-participant level. This modelled the onsets of picture stimuli, emotional judgement task, and movement parameters against BOLD activity pre- and post-intervention. We used Helmert contrast codes for t-tests comparing: dehumanized vs humanized targets [3:1], homeless vs humanized targets [6:1], and homeless vs drug targets [1:1]. Due to an error in the coding of stimuli presentation, 2 trials per participant were modelled and partitioned out of any other further analyses. We generated second-level F contrast [1] images for group level statistics using the first-level t-test images.

## Results

### Behavioral Data

#### Emotional Judgment Task

152 emotional judgement ratings were excluded from analysis due to logging errors or lack of responses. We analyzed data separately for each scanning session and in relation to each participant’s proportion of judgements within each target (percentage of trials the participant rated with pride, envy, pity, or disgust). We conducted post-hoc analyses in accordance with the multiple regression approach to analyzing contingency tables (Beasley, & Schumacker, 1995), and a Bonferroni corrected threshold of *p* = .0015625 was used.

Our chi-square test of independence revealed a significant relationship between the target images and subsequent responses in the pre-intervention condition (χ^2^ (*21*, *N* = 1262) = 737.41, *p* <.001). Post-hoc analyses revealed at least 1 significantly disproportionate emotional judgement response for all pre-intervention targets (Table 1). Consistent with the prior literature, drug-addicted targets were consistently rated with feelings of disgust (57.05%, *p* <.00001). However, the proportion of disgust ratings for homeless targets was not significant (21.34%, *p* = .03), but pity ratings were significantly the most common response (67.68%, *p* <.00001).

A chi-square test of independence revealed a significant relationship between the target images and subsequent emotional judgement responses in the post-intervention condition (χ^2^ (*21*, *N* = 1168) = 781.74, *p* <.001). Post-hoc analyses revealed at least 1 significantly disproportionate emotional judgement response for all targets post-intervention (Table 1). Disgust ratings for the homeless remained non-significant across scanning sessions. Envy and pity responses for the homeless increased post-intervention, whilst pride responses decreased. The rich group saw the most dramatic change in response (Figure 10).

**Table 1.**
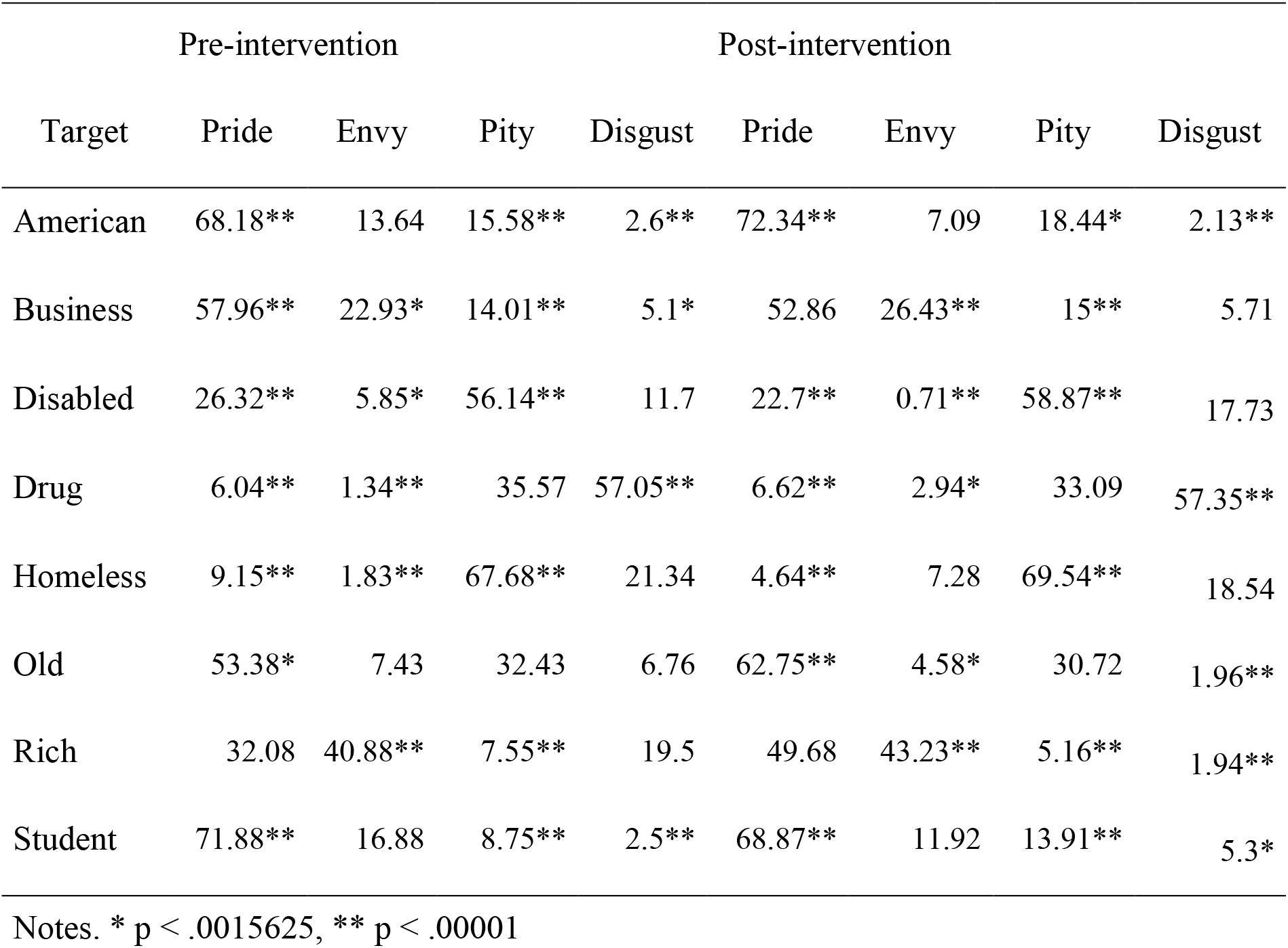
Mean proportion of emotional judgments (%) per target per scanning session.

**Figure 10.**
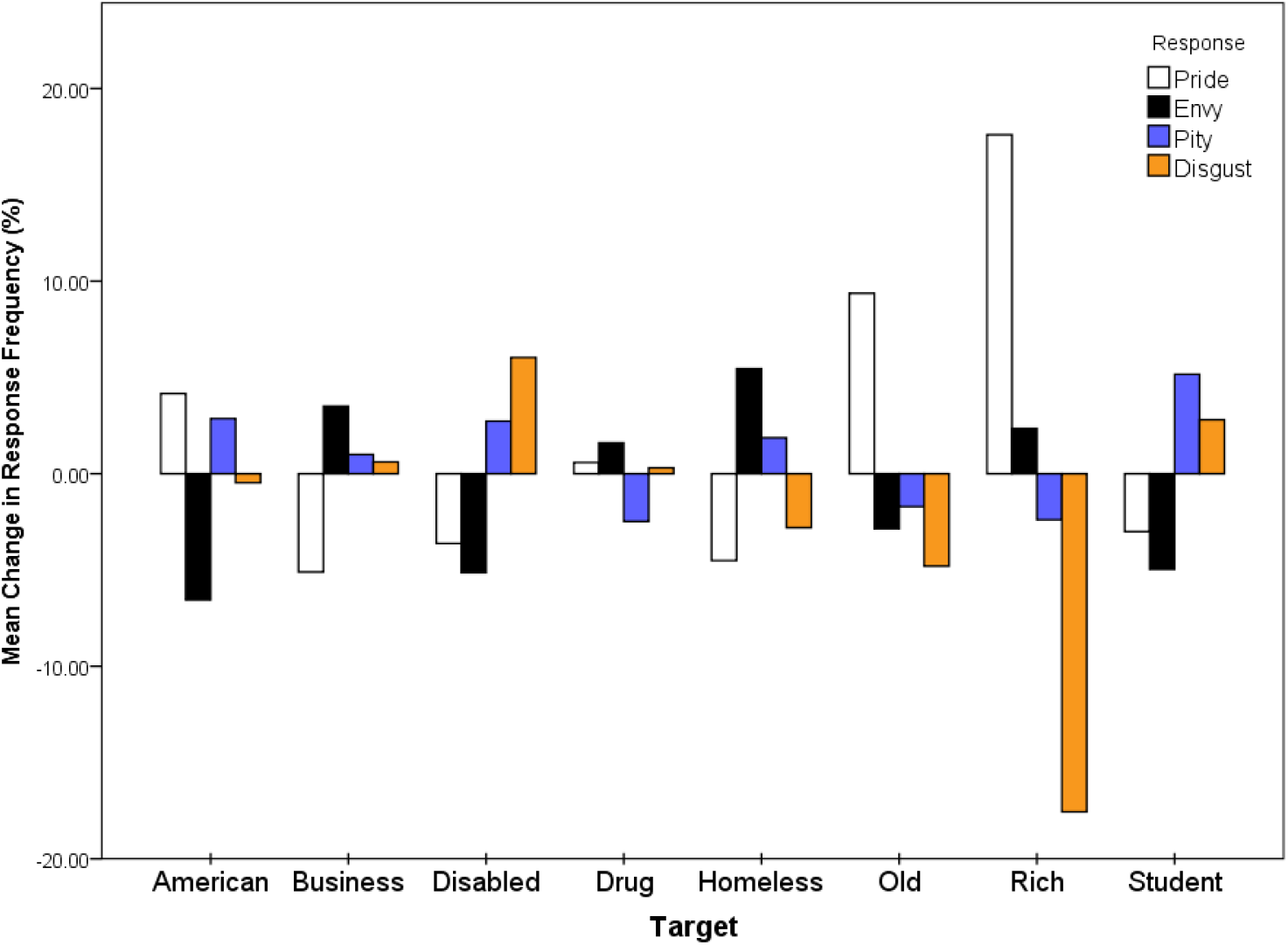
Average change in responses (%) following the intervention.

**Figure 11.**
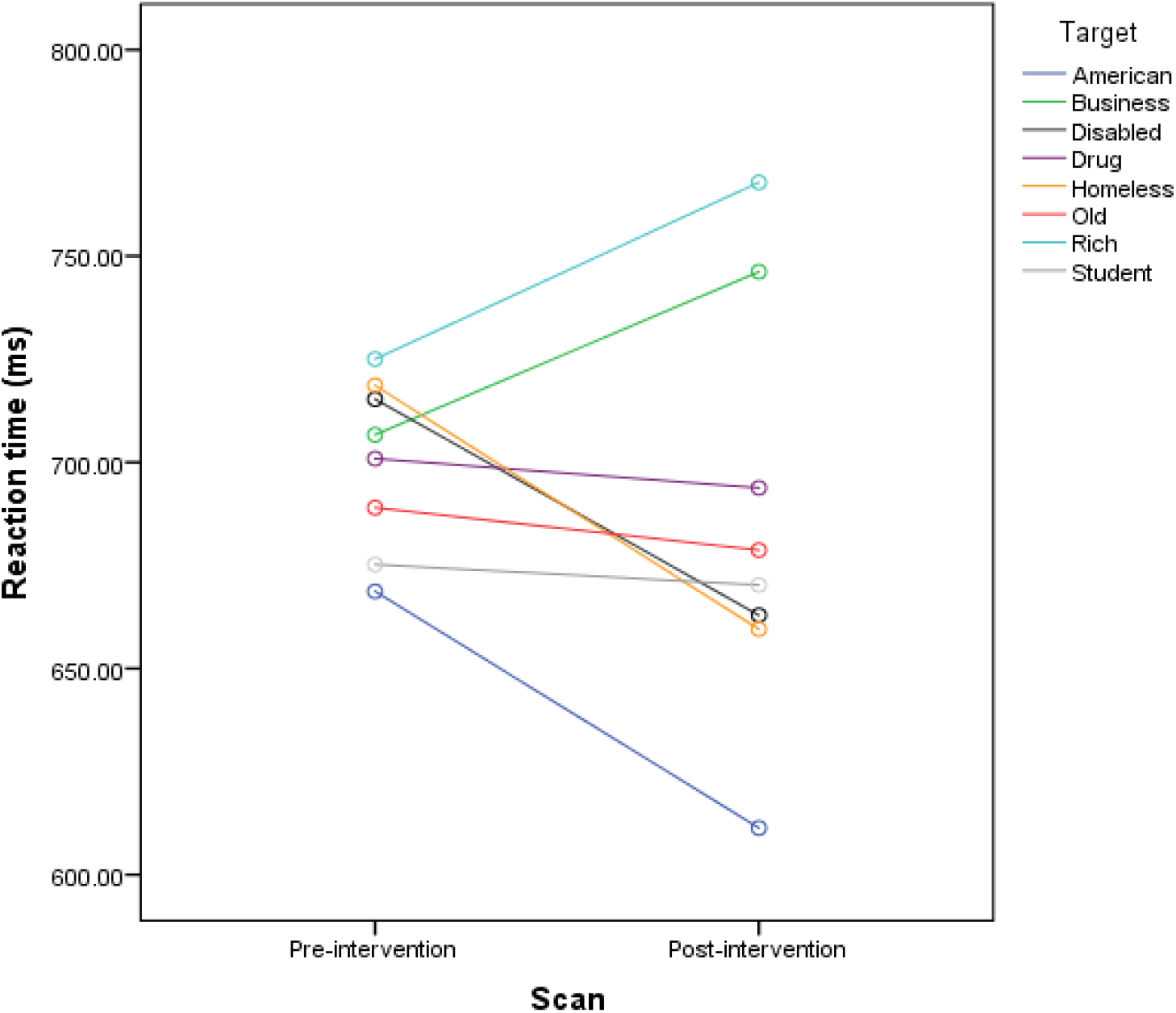
Mean reaction times (ms) to target across scanning sessions.

### Reaction Times

We excluded 2 participants from reaction time analyses and 77 trials due to logging errors, lack of responses, or outliers (+/-2.5 SD). We collapsed the data into mean reaction times per target for each scanning session. Mauchly’s test suggested the assumption of sphericity was not violated (χ^2^ (27) = 27.414, *p* = .577). A repeated measures ANOVA revealed a significant main effect of target on reaction time (*F* (7, 56) = 2.55, *p* < .05, *η_p_^2^*= .24), though a scan x target interaction was not found (*F* (7, 56) = 2.55, *p* < .05, *η_p_^2^*= .24). Pairwise comparisons with Fisher’s least significant difference correction revealed significant differences in reaction time between American (*M* = 639.11ms, *SD* = 35.35) and Rich (*M* = 751.78ms, *SD* = 47.47) targets (*p* <.05), and American and Business (*M* = 720.57ms, *SD* = 52.09) targets (*p* <.05). When using a stricter threshold (bonferroni correction), no significant differences were seen post-hoc.

### fMRI Data

#### Whole-Brain Contrasts

Whole-brain contrasts at the family-wise error (FWE) corrected threshold of *p* < .05 yielded minimal results. Due to the potentially underpowered nature of the study, we lowered the threshold to an uncorrected *p* < .001 for exploratory purposes. The reported results are for peak-values at this threshold unless otherwise stated. All coordinates stated are in accordance with MNI space. We labelled corresponding neuroanatomical locations via cross-referencing the SPM Anatomy Toolbox (Eickhoff et al. 2005), SPM’s built-in neuromorophometric labelling, and NeuroSynth (Yarkoni, Poldrack, Nichols, Van Essen, & Wagner, 2011).

##### Dehumanized vs Humanized Targets

A whole-brain F contrast of dehumanized vs humanized targets pre-intervention showed 1 cluster of 1 voxel at the FWE corrected *p* <.05 threshold in the insula (Figure 12). Peak-level activation was significantly greater for dehumanized targets in an anterior region of the right insula (*F* (1, 10) = 160.01, *p* < .01, at x = 33, y = 11, z = 8). Peak-level voxels at the uncorrected threshold also included reduced activity in the cuneus (*F* (1, 10) = 45.14, *p* < .001, at x = 0, y = −85, z = 26), precuneus (*F* (1, 10) = 39.08, *p* < .001, at x = 3, y = −61, z = 32), and the left inferior parietal lobule (*F* (1, 10), *p* < .001, x = −39, y= −55, z = 50).

**Figure 12.**
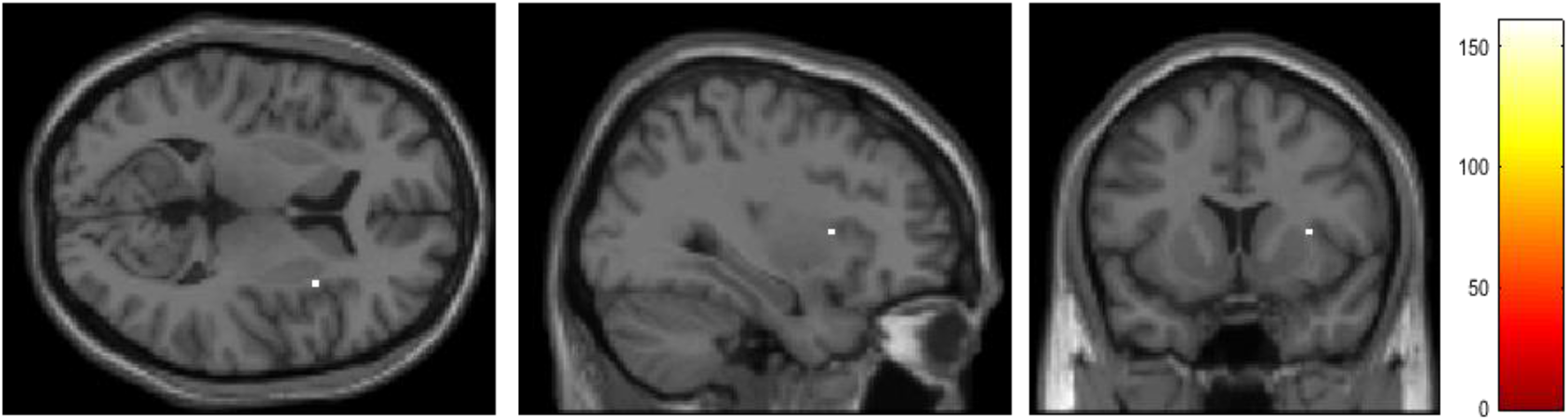
Single voxel cluster in right insula foe Dehumanized> Humanized targets pre-invention (FEW *p*<0.5 threshold).

A whole-brain F contrast of dehumanized vs humanized targets post-intervention did not reveal the same pattern of activity as the prior scanning session. Instead, activations were in regions such as the left fusiform gyrus (*F* (1, 10) = 37.26, *p* < .001, at x = −27, y = −49, z = −10; results in Table 2).

**Table 2.**
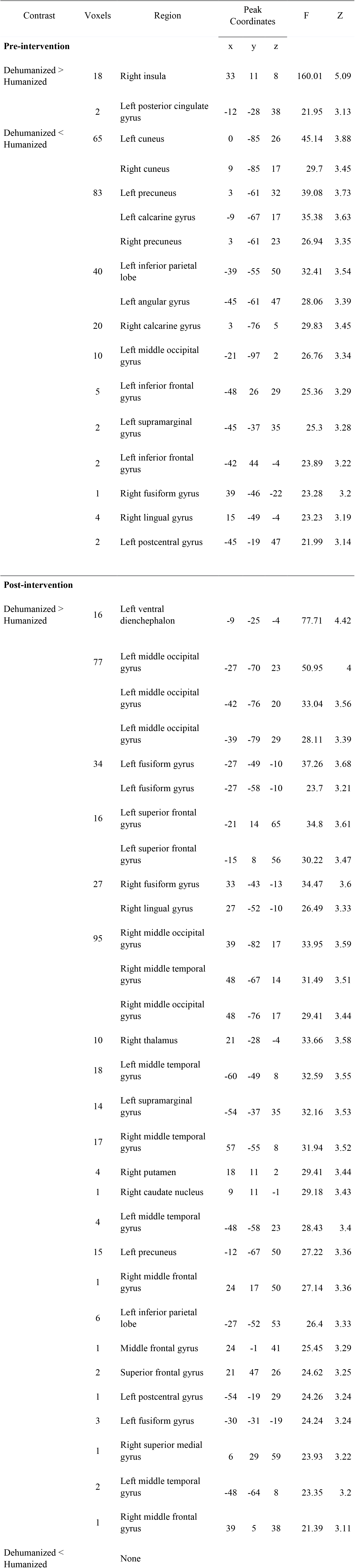
Main effects contrasts for Dehumanized vs Humanized targets. All values reported are peak-level voxels at the *p* < .001 (uncorrected) 0 voxel extent threshold.

#### Homeless vs Humanized targets

Pre-intervention, no significant activity was greater for homeless over humanized targets. The greatest differential peak-level activation was observed in the left middle temporal gyrus (*F* (1, 10) = 3.85, *p*<.001, at x = −54, y = −64, z = −4), demonstrating less activity for the homeless. Other regions showing reduced activity included the left inferior parietal lobe (*F* (1, 10) = 27.95, *p* < .001, at x = −45, y = −34, z = 35), right fusiform gyrus (*F* (1, 10 = 23.82, *p* < .001, at x = 42, y = −46, z = −16), and subgenual anterior cingulate cortex (*F* (1, 10 = 23.45, *p* < .001, at x = −3, y = 29, z = −1). Post-intervention, we did not observe the same pattern of activity (Table 3). Significantly greater activity in a different region of the right fusiform gyrus was present (*F* (1, 10) = 45.91, *p* < .001, at x = 33, y = −40, z = −10). The greatest differential peak-level activation was observed within a region of occipital gyri (*F* (1, 10) = 85.70, *p* < .001, at x = −42, y = −70, z = 29).

**Table 3.**
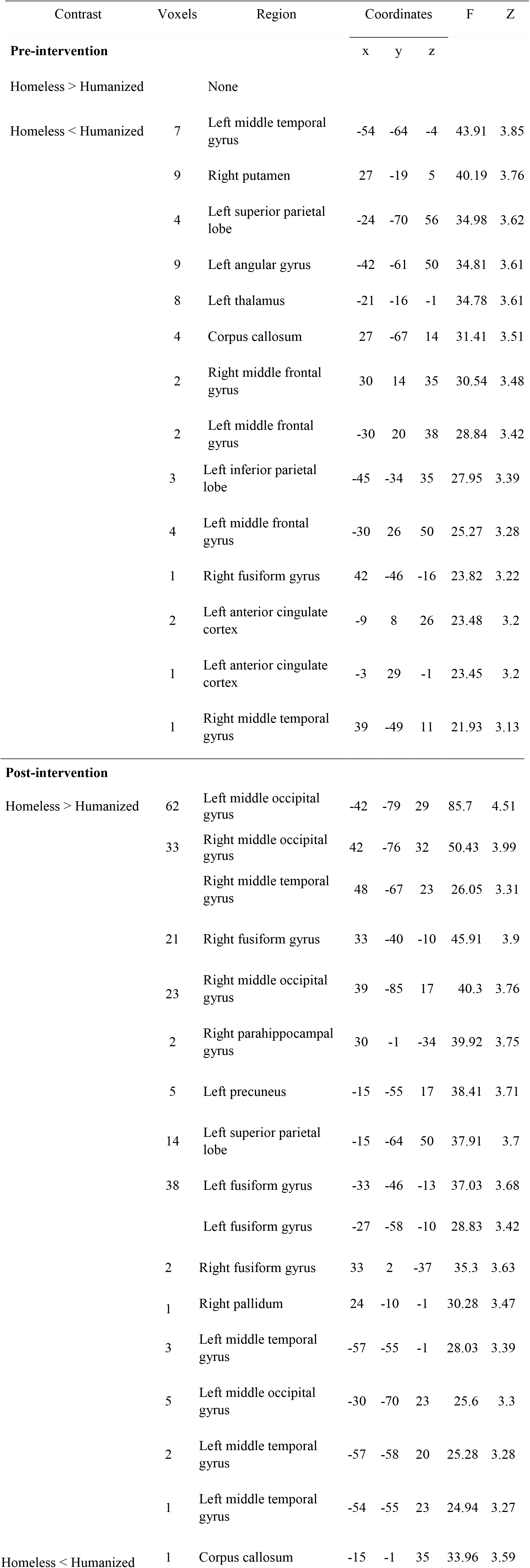
Main effects contrasts for Homeless vs Humanized targets. All values reported are peak-level voxels at the *p* < .001 (uncorrected) 0 voxel extent threshold.

#### Homeless vs Drug-addicted Targets

For pre-intervention scans, a region of the right anterior insula showed the largest increase in activation for drug-addicted targets (*F* (1, 10) = 42,67, *p* < .001, at x = 39, y = 5, z = −4). Significant activations also included the left inferior parietal lobe (*F* (1, 10) = 42,42, *p* < .001, at x = −57, y = −22, z = 44), and left midcingulate gyrus (*F* (1, 10) = 28.96, *p* < .001, at x = −3, y = 2, z = 32). Post-intervention, we did not see differential insula response and significant activation was observed in both a superior (*F* (1, 10 = 102.52, *p* < .001, at x = 33, y = −43, z = 56) and inferior region (*F* (1, 10) = 56.18, *p* < .001, at x = −39, y = −40, z = 50) of the left inferior parietal lobe.

**Table 4.**
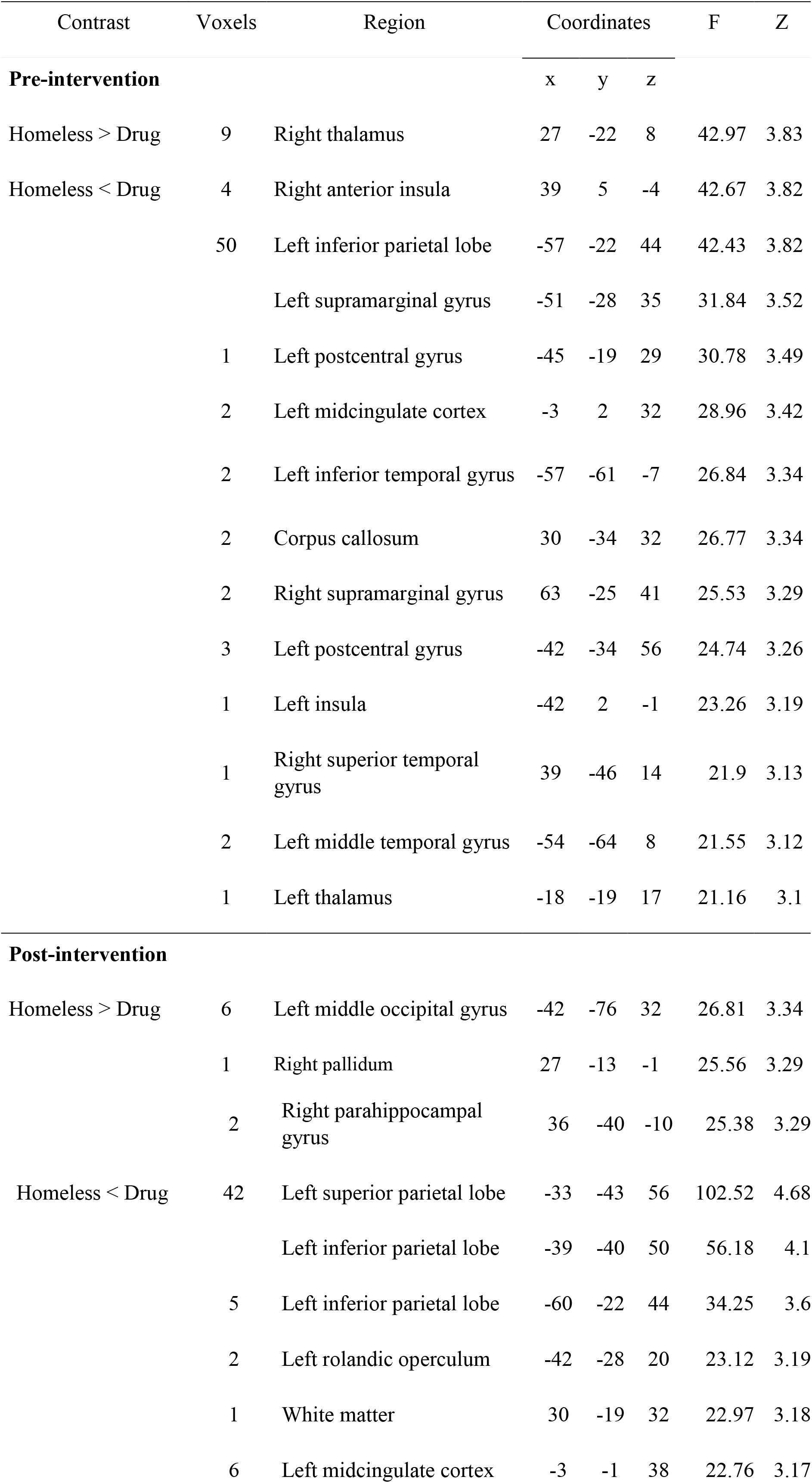
Main effects contrasts for Homeless vs Drug-addicted targets. All values reported are peak-level voxels at the p < .001 (uncorrected) 0 voxel extent threshold.

### Regions of Interest (ROI) Analysis

Based on our hypotheses, and the significant peak-value voxels in pre-intervention contrasts, we extracted beta-weights from three coordinates in the homeless vs humanized contrast. The first region was the right insula (x = 33, y = 11, z = 8), which demonstrated greater activation for dehumanized than humanized targets. The second two were the left IPL (x = −45, y= −34, z = 35) and left middle temporal gyrus (x = −54, y = −64, z = −4) due to their proximity within the social cognition network. To test whether effects were unique to the homeless, or the whole low-low group of the SCM, we ran ROI analysis on the homeless vs drug-addicted contrast images.

#### Right Insula

In the homeless vs humanized contrast, beta values for the peak voxel in the right insula demonstrated greater activity for homeless targets (*M* = 2.51, *SD* = 3.23), and this was reduced post-intervention (*M* = 1.18, *SD* = 6.34). A repeated-measures ANOVA failed to find a significant difference in right insula betas between scanning sessions (*F* (1, 10) = .693, *p* = .42, *ηp^2^* = .07). Homeless targets evoked marginally less activity than drug-addicted targets pre-intervention (*M* = −.89, *SD* = 1.53), and post-intervention (*M* = −.63, *SD* = 1.30). A repeated measures ANOVA found no significant difference for this contrast (*F* (1,10) = .29, *p* = .60, *ηp^2^* = .03).

#### Left Inferior Parietal Lobe (IPL)

In the homeless vs humanized contrast, beta values for the peak voxel in the left IPL demonstrated decreased activity for homeless targets pre-intervention (*M* = −4.61, *SD* = 2.89), and this difference was minimized post-intervention (*M* = −.71, *SD* = 5.59). A repeated measures ANOVA found this to be a significant effect (*F* (1, 10) = 8.83, *p* = .01, *ηp^2^* = .47). A repeated measures ANOVA for the homeless vs drug contrast found no significant change in activation discrepancy between targets was observed following the intervention (*M* = −1.37, *SD* = 1.45) or two (*M* = −1.30, *SD* = 2.00; *F* (1, 10) = .015, *p* **=**, *ηp^2^* = .00).

#### Left Middle Temporal Gyrus (MTG)

In the homeless vs humanized contrast, activity in the left MTG was decreased for homeless targets (*M* = −6.17, *SD* = 3.09), and this changed to an increased activation post-intervention (*M* = 7.54, *SD* = 10.91). A repeated measures ANOVA found this to be a significant effect (*F* (1, 10) = 8.83, *p* < .05, *ηp^2^* = .47). Whilst left MTG activity was decreased for homeless over drug targets pre-intervention (*M* = −5.01, *SD* = 4.56), this difference was reduced post-intervention (*M* = −1.25, *SD* = 2.35). A repeated measures ANOVA found this change in activation discrepancy to be significant (*F* (1, 10) = *p* < .05 *ηp^2^* = .45).

### Summary of results

Participants consistently rated drug-addicted targets with feelings of disgust across both sessions, but the homeless with feelings of pity. Reaction time data revealed no significant differences for the homeless and drug-addicted targets. One region showed significant activation at the family wise error corrected threshold for dehumanized targets, the right insula – pre-intervention, but not post-intervention. The increase between scanning sessions was not significant following regions of interest analysis. No significant difference in mPFC was apparent response between dehumanized and humanized targets pre-intervention. Two other regions in the social cognition network of particular interest, the left IPL and MTG, demonstrated significantly decreased activity for homeless targets in the pre-intervention condition. ROI analysis for these locations demonstrated a significant increase in BOLD following contact. The change in the MTG appeared to be unique to the homeless, whereas a change in left IPL activity appeared to occur alongside drug-addicted targets.

## General Discussion

This two-part study primarily aimed to see whether dehumanization effects, as indexed by activity in the social cognition network, could be altered following contact with an extreme outgroup, the homeless. We found mixed findings in relation to both dehumanization effects towards the homeless, and brain response following a social contact intervention to diminish this bias. The results from experiment 1 replicate previous dehumanization effects (Harris and Fiske, 2006), as seen by lessened activity in the mPFC when viewing images of homeless individuals. Although we saw no significant effects of contact, time spent in the intervention significantly predicted change in mPFC response following social contact. In experiment 2, one region showed significant activation at the FWE corrected threshold for dehumanized targets, the right insula (pre-intervention, but not post-intervention). After easing our statistical threshold, results did not show a significant reduction in mPFC activity, but did in other areas of the social cognition brain network. ROI analyses of the left IPL and MTG showed an increase in activity following the contact intervention, but no change in right insula signal.

### Dehumanization

The two experiments did not reach agreement on the role of the mPFC. Considering the strength of the prior literature, and the low-sample size used here, concluding that no dehumanization effects in the mPFC exist would be unjustified. The findings in the first study demonstrated that change in mPFC to the homeless altered as a product of time spent in the intervention. As this time was variable, perhaps the lack of a main effect was due to participant’s different responses to the interventions. Furthermore, to participate in the study, subjects had to commit to both scanning sessions, and the intervention, knowing they would hold an interaction with homeless individuals. If engaging with the homeless is often avoided due to anticipated exhaustion (Cameron, Harris, and Payne, 2016), those who were willing to volunteer may not have anticipated said tiring, and were more likely to engage social cognition for the homeless prior to the study. However, if this were just for homeless targets and not the low-low SCM group as a whole, we would have seen decreased mPFC activity for drug-addicted targets – this was not the case. Feasibly, the recruitment bias could have meant the sample also engaged social cognition for the drug-addicted, due to the low salience of intergroup borders. Where this becomes inconsistent with the literature however, is in relation to the hypothesized affective processes.

### Affective change

While we observed no difference in amygdala response toward the homeless, the second experiment demonstrated increased insula activity for dehumanized targets pre-intervention, but not post-intervention. These findings seem consistent with previous studies (Harris & Fiske, 2007) suggesting that both homeless and drug-addicted targets should evoke increased insula activity as a product of disgust. Yet, when contrasting homeless and drug-addicted targets, an anterior region of the insula revealed differential activation. Despite both belonging to the low-low SCM category, drug-addicted targets evoked significantly more activity than homeless targets. Previous results (Harris & Fiske, 2011) have found right anterior insula activity negatively correlates with perceived warmth. Additionally, our behavioral data showed participants consistently rated the homeless with feelings of pity, but the drug-addicted with disgust. Thus, participants in the present study may be viewing homeless targets as incompetent, but not cold or ill intending.

Given that participants showed brain and behavioral disgust responses to the drug-addicted, the expectation would be a subsequent downregulation of mPFC for the drug-addicted – this was not the case. This does not completely disregard schema-triggered affect as having an impact on humanized perception. Instead, this highlights: (1) the role of affect as an indirect mediating variable, rather than a definitive moderating factor; and (2) a network response, and not just the mPFC, indexes dehumanization (Harris, & Fiske, 2006; 2007).

### Rehumanization

The pre-intervention homeless-humanized contrast (experiment 2) demonstrated significantly decreased activity in the left MTG and IPL for homeless targets. Regions of interest analysis demonstrated a significant increase in BOLD for these regions following contact. Previous data has shown modulation of these two regions as product of motivation to infer mental states in dehumanized targets (Harris & Fiske, 2007). Moreover, research conducted subsequent to the present study (Bruneau, Jacoby, Kteily, & Saxe, 2018) has clarified the unique role of the left IPL in dehumanized perception beyond like/dislike of social groups. In addition, social cognition tasks outside of dehumanized contexts have consistently shown activation in these regions (Arora et al., 2015; Van der Meer, Groenewold, Nolen, Pijenborg, & Aleman, 2011; Van Overwalle 2009; Blakemore, 2012). Overall, the data shows altered social cognition brain network response to homeless targets following contact, and this seemed to occur regardless of the participants’ affective responses. Although our statistical threshold and circular analysis does not allow for detailing the specific mechanisms of change, the ROI analysis lends some support for contact as an intervention for humanizing extreme out-groups. In light of these results, we have several suggestions for refining methodology.

### Future research

Our results indicate the homeless did not elicit frequent feelings of disgust. Importantly, this pushes for a re-evaluation of how the SCM conceptualizes its categories, and emotional responses, but could potentially be a reflection of social desirability effects. We made use of an explicit forced-choice paradigm for emotional judgements. However, to lend greater insights into perceivers’ stereotype contents, a richer measure of affect may be required. It would also be intriguing to see the implementation of advanced fMRI statistics, such as effective connectivity analysis, to elucidate the exact role of differences in schema-triggered affect and dehumanization effects.

Though the results presented here do warrant further investigations of affective influence, we have presented a justified case against schema-triggered affect being the sole, dominant factor attenuating dehumanized perception. Studies show that prior motivation seems to be a driving force behind dehumanization (Cameron et al., 2016), yet behavioral correlates of this motivation in neuroimaging studies remains unmeasured. If motivation is a causal factor, it is imperative future studies include this in design. Beta-weights extracted from the current data demonstrated the variable effects of contact as a product of time in the interaction. By including measures of motivation and anticipated exhaustion as a factor in fMRI analyses, we can start to understand the neurocognitive mechanisms influencing contact’s efficacy.

Lastly, the literature has made some commentary concerning secondary effects (see Cloutier, Li, and Correll, 2013), but has not addressed, or tested this directly. For instance, our second experiment showed the left IPL increased for both homeless and drug-addicted targets following contact. One would not expect this effect if implicit attitudes towards social groups remained mutually exclusive. A posteriori explanations would be speculative, and are not due for these cross-group effects, but the data is certainly worth noting. These findings open potential avenues of research, especially if researchers contrast the dehumanized groups with those on relevant social spectra (e.g. socioeconomic status) such as the rich. Future studies could attend to the dynamic, intertwined relationship between attitudes across social groups.

## Conclusion

The present study yielded mixed results. The first experiment replicated previous dehumanization effects, as seen by decreased mPFC activity to homeless targets. Additionally, mPFC activity seemed to change as a product of time spent in the contact intervention. The second experiment did not show this mPFC response. Rather, results of interest included a wider array of regions previously indicated in dehumanized perception: Insula, left IPL, and left MTG. Altered activity in the latter two regions lends support to the use of contact as a practical intervention. Additionally, we did not see the homeless rated with feelings of disgusted, and this was consistent with the discrepancy in Insula activation between homeless and drug-addicted targets. However, Insula activity did not seem to change alongside the left IPL or left MTG, pulling into question the role of schema-triggered affect in dehumanization. Although these results lend some support to the use of contact as an intervention, the cognitive mechanisms underlying this process need further delineating.

## Acknowledgements

Appreciation goes to the faculty at University College London and the Duke-UNC Brain Imaging & Analysis Center. Many thanks should also go to the staff at Urban Ministries of Durham soup kitchen, and the homeless individuals who formed a crucial part of the contact intervention. Lastly, a special thanks to Dr. Samuel Evans and Dr. Tae Twomey for their advice on SPM.

